# PGC-1β prevents statin-associated myotoxicity in oxidative skeletal muscle

**DOI:** 10.1101/218636

**Authors:** François Singh, Joffrey Zoll, Urs Duthaler, Anne-Laure Charles, Gilles Laverny, Daniel Metzger, Bernard Geny, Stephan Krähenbühl, Jamal Bouitbir

**Affiliations:** Department of Clinical Pharmacology and Toxicology, Department of Biomedicine, University Hospital, Hebelstrasse 20, 4031 Basel, Switzerland; Université de Strasbourg, Fédération de Médecine Translationelle, Equipe d’Accueil 3072, Institut de Physiologie, Faculté de Médecine, 11 rue Humann, 67085 Strasbourg, France; Swiss Centre for Applied Human Research (SCAHT), Switzerland; Institut de Génétique et de Biologie Moléculaire et Cellulaire, Centre National de la Recherche Scientifique UMR7104, Institut National de la Santé et de la Recherche Médicale U964, Université de Strasbourg, Strasbourg, France

**Keywords:** atorvastatin, myopathy, PGC-1β, apoptosis, reactive oxygen species (ROS), mitochondrial proliferation

## Abstract

Statins are generally well-tolerated, but can induce myopathy. Statins are associated with impaired expression of PGC-1β in human and rat skeletal muscle. The current study was performed to investigate the relation between PGC-1β expression and function and statin-associated myopathy. In WT mice, atorvastatin impaired mitochondrial function in glycolytic, but not in oxidative muscle. In PGC-1β KO mice, atorvastatin induced a shift from oxidative type IIA to glycolytic type IIB myofibers mainly in oxidative muscle and mitochondrial dysfunction was observed in both muscle types. In glycolytic muscle of WT and KO mice and in oxidative muscle of KO mice, atorvastatin suppressed mitochondrial proliferation and oxidative defense, leading to apoptosis. In contrast, mitochondrial function was maintained or improved and apoptosis decreased by atorvastatin in oxidative muscle of WT mice. In conclusion, PGC-1β has an important role in preventing damage to oxidative muscle in the presence of a mitochondrial toxicant such as atorvastatin.

---

HMG-CoA reductase inhibitors (statins) are effective in lowering of LDL-cholesterol with clinically relevant beneficial effects in the prevention of cardiovascular events (1,2). Muscle toxicity is one of the most common adverse event encountered in patients treated with statins, with up to a quarter of patients affected (3). Statin-associated muscle toxicity ranges from asymptomatic elevation of creatine kinase (CK) activity to myalgia with or without increased CK activity and finally to rhabdomyolysis (4), which can be life-threatening (5). Several mechanisms leading to myopathy have been proposed, among them dysregulated muscle Ca^2+^ homeostasis (6,7), proteolysis due to increased atrogin-1 expression (8,9), oxidative stress (10), decreased skeletal muscle coenzyme Q_10_ concentrations (7) and inhibition of complex III of the respiratory chain (11). Impairment of mitochondrial function triggering apoptosis appears to be a central factor in many of the proposed mechanisms (12–15). Moreover, statins have been reported to induce apoptosis in skeletal muscle (10,13). Recently, we showed in humans and in rats that statins can trigger apoptosis in fast glycolytic skeletal muscle, and that this mechanism is ROS dependent (13). In comparison, oxidative skeletal muscle, which has a high mitochondrial content, was not affected by statins. This is in agreement with experiments in L_6_ myoblasts, where stimulation of mitochondrial hormesis through PGC-1β was partially protective against the harmful effects of statins (14). Glycolytic muscle appears to be more sensitive to statins than oxidative muscle, and a high mitochondrial content appears to be protective.

Interestingly, long-term treatment with statins has been reported to decrease mitochondrial content in skeletal muscle, underscoring mitochondrial toxicity of statins (16–18). The members of the peroxisome proliferator-activated receptor γ co-activator 1 family (PGC-1) are considered as key regulators of mitochondrial biogenesis, content and function (19–21). Our previous data showed that statins can protect mitochondria in the highly oxidative cardiac muscle by triggering mitochondrial hormesis via stimulation of PGC-1β expression in both humans and rats (22). On the other hand, in glycolytic skeletal muscle, we observed a repression of mitochondrial proliferation by statins correlated to impaired expression of PGC-1α and PGC-1β. This mechanism, termed mitochondrial hormesis, proposes that low concentrations of mitochondrial ROS can trigger mitochondrial biogenesis and thereby counteract oxidative stress and re-establish homeostasis (23–26).

These observations suggested a role of PGC-1β in statin-associated muscle pathology. While regulation and function of PGC-1α in skeletal muscle is well established (27), the role of PGC-1β is currently less clear and debated in the literature (28–30). Recently, we described a mouse model with a selective ablation of PGC-1β in skeletal muscle fibers at adulthood (PGC-1β^(i)skm-/-^ mice) (31), and showed that PGC-1β not only coordinates the expression of genes controlling mitochondrial structure and oxidative phosphorylation, but also genes involved in antioxidative defense such as mitochondrial superoxide dismutase 2 (SOD2).

Since statins are associated with mitochondrial damage including impairment of oxidative phosphorylation and mitochondrial proliferation as well as increased mitochondrial ROS production, we speculated that myotoxicity of statins could be increased in PGC-1β^(i)skm-/-^ compared to wild-type mice, suggesting a preventive role of PGC-1β in statin-associated myotoxicity. Our current study provides evidence that not only antioxidative capacity and mitochondrial content are important factors for the protection from statin-associated myotoxicity, but also the ability to trigger mitochondrial adaptive pathways such as mitochondrial proliferation, which is at least partially dependent on the function of PGC-1β.

## Results

### In wild-type and PGC-1β^(i)skm-/-^ mice, atorvastatin treatment impaired skeletal muscle function

To confirm the efficiency of the treatment, we first analyzed the plasma atorvastatin concentration by LC-MS/MS (Figure 1A). As no differences in the plasma atorvastatin concentration were observed between WT-ATO and KO-ATO groups (data not shown), we pooled the values of these two groups. The plasma concentration reached a mean of 20 nM (range 9 nM to 33 nM) in statin-treated animals, which is comparable to what is observed in patient plasma (32) and is in the range of the IC_50_ for the inhibition of HMG-CoA reductase (33). In the plasma of control groups (WT-CTL and KO-CTL), atorvastatin was not detectable (data not shown). Plasma creatine kinase activity reflects muscle damage and can be used as a biomarker for the detection of drug-induced myopathies. Atorvastatin treatment increased the plasma creatine kinase activity in both WT and KO mice (two-way ANOVA treatment main effect p=0.0008, F=17.65), reaching statistical significance for KO-ATO compared to KO-CTL mice (Figure 1B). This finding suggested the existence of atorvastatin-associated muscular alterations, particularly in PGC-1β^(i)skm-/-^ mice. To confirm this on a functional level, we performed grip strength tests, and observed that statin treatment decreased mice peak force in both groups compared to their respective control group (two-way ANOVA treatment main effect p=0.0027, F=10.25) (Figure 1C). No differences were found in the body weight at the end of the protocol in any group (Figure 1D), suggesting that the decrease in muscle strength was associated with muscle dysfunction and not with a general effect on the animals.

**Figure 1:**
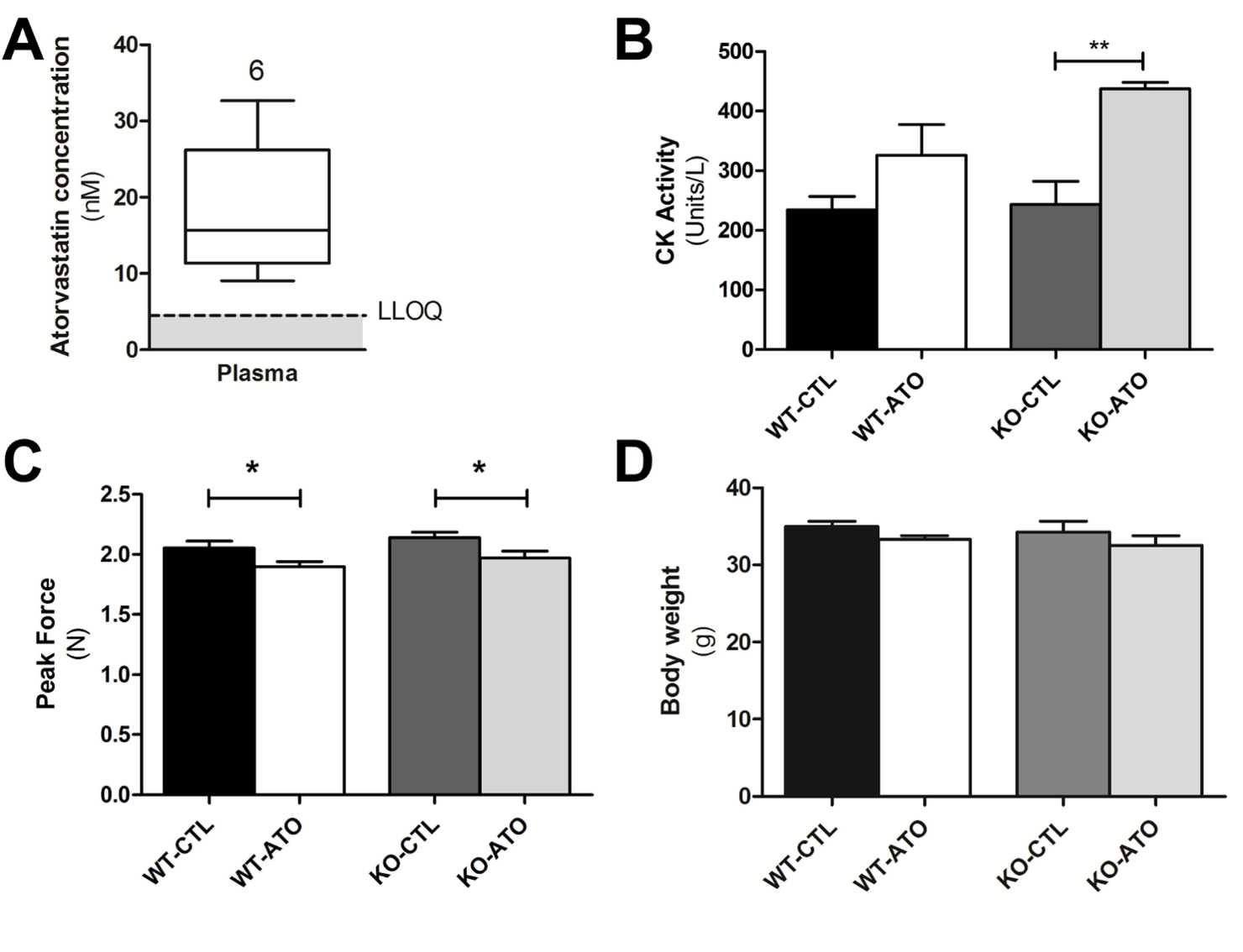
Atorvastatin treatment impaired skeletal muscle physiology. **(A)** Plasmatic atorvastatin concentration. The upper and lower limits of the boxes correspond to the interquartile range and the middle line is the median of the data set. The limits of the whiskers are the minimum and maximum values. Linear regression through the mean value of the boxes was established. **(B)** Plasmatic creatine kinase activity. **(C)** Forelimb and hindlimb peak force. **(D)** Mice body weight at the end of the study. **(B-D)** Results are expressed as mean ±SEM; n=8−13 per group, *p<0.05, **p<0.01.

### Atorvastatin treatment induced a switch from oxidative to glycolytic muscle fiber phenotype

To get a better insight into the suspected muscular alterations, we performed a hematoxylin and eosin staining on sections of two metabolically different hindlimb muscles: the superficial gastrocnemius, which is mostly glycolytic, and the vastus intermedius, which is mostly oxidative. We observed a decrease of fiber area in the glycolytic muscle (Figure S1A) of both WT-ATO and KO-ATO mice compared to their respective control group (two-way ANOVA treatment main effect p=0.0027, F=10.25). Interestingly, atorvastatin decreased the fiber area also in the oxidative muscle of KO mice, but not of WT mice (two-way ANOVA interaction effect p=0.0009, F=21.51) (Figure S1B).

To obtain a better insight of the relation between fiber type and statin toxicity, we determined myosin heavy chain isoform (MHC) I, IIA, IIX, and IIB expression in both muscle types. In the glycolytic gastrocnemius muscle, both groups of mice treated with statins exhibited a switch from IIA myofibers, (two-way ANOVA treatment main effect p=0.0002, F=37.10) which are characterized by mitochondrial abundance and oxidative metabolism (34), and IIAX myofibers (two-way ANOVA treatment effect p=0.0284, F=6.80) towards IIB glycolytic myofibers (two-way ANOVA treatment main effect p=0.0004, F=30.28) (Figure 2A). In comparison, in the oxidative vastus intermedius muscle of WT mice, statins did not affect fiber type composition (Figure 2B). However, in the oxidative muscle of KO mice, atorvastatin induced a massive switch from fast-twitch oxidative IIA (two-way ANOVA interaction effect p=0.0010, F=29.31), IIAX (two-way ANOVA interaction effect p=0.0243, F=8.95), and IIX (two-way ANOVA interaction effect p=0.0106, F=13.38) towards IIB glycolytic myofibers (two-way ANOVA interaction effect p=0.0002, F=66.90).

**Figure 2:**
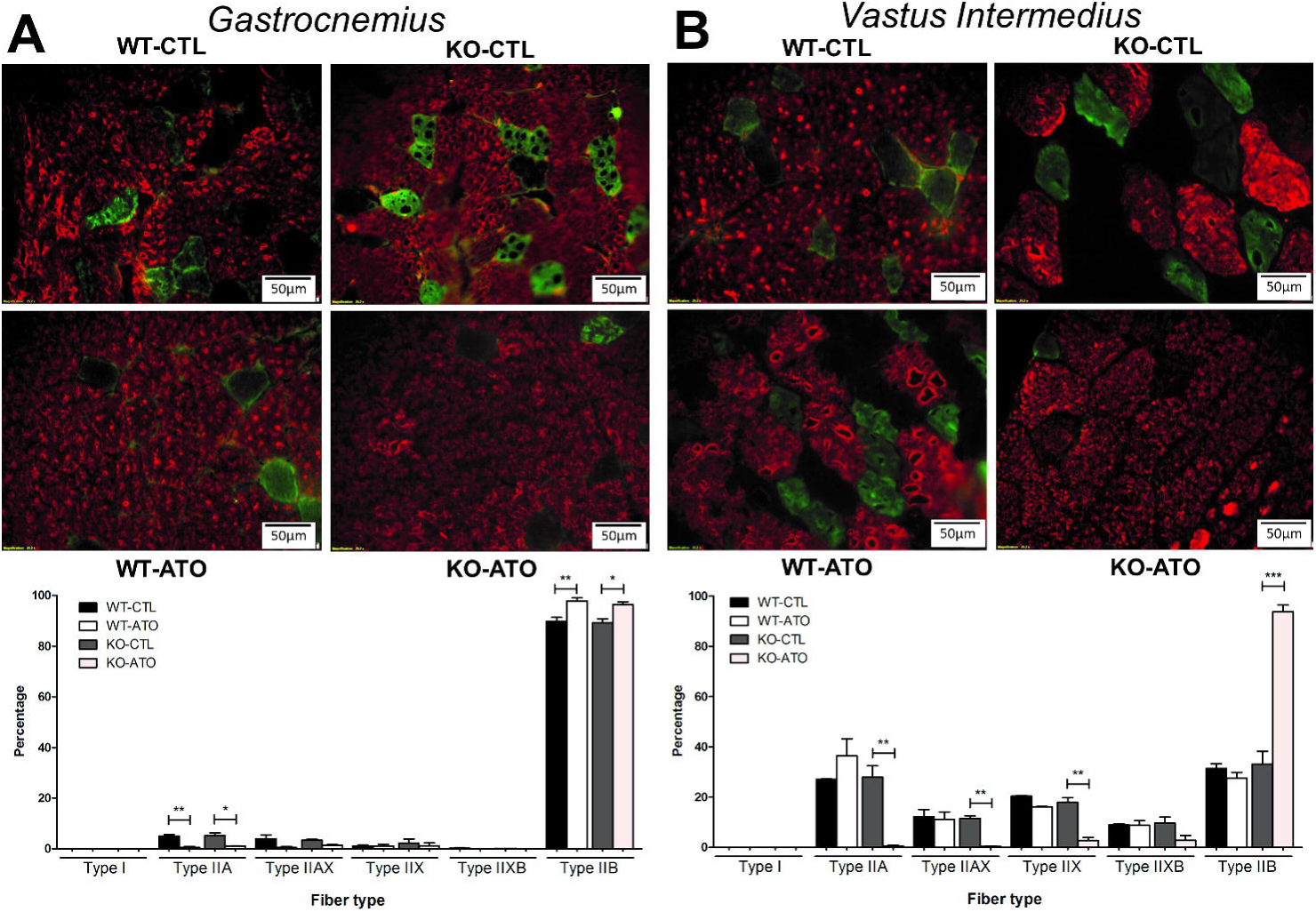
Atorvastatin treatment induced a switch in fiber type in the glycolytic skeletal muscle and in the oxidative skeletal muscle of PGC-1β deficient mice. **(A)** Fiber typing of superficial gastrocnemius fiber sections and quantification. **(B)** Fiber typing of vastus intermedius fiber sections and quantifications. **(A-B)** Shown are type I (blue), type IIA (green), type IIAX (intermediate green), type IIX (dark), type IIXB (intermediate red), and type IIB (red). Results are expressed as mean ±SEM; n=8−13 per group, *p<0.05, **p<0.01, ***p<0.001.

Taken together, these results suggested that the switch from the oxidative to the glycolytic phenotype potentially could increase the susceptibility of oxidative fibers to atorvastatin. Since statins are known to impair mitochondrial function in glycolytic skeletal muscle (13,22), it was of interest to study mitochondrial function and mitochondrial adaptations in PGC-1β^(i)skm-/-^ mice.

### Atorvastatin impaired mitochondrial function in glycolytic muscle of wild type and PGC-1β^(i)skm-/-^ mice whereas oxidative muscle exhibited defects only in PGC-1β ablated mice

In agreement with previous results obtained in the glycolytic tibialis anterior muscle (31), histochemical staining revealed decreased activity of mitochondrial complex I (NADH dehydrogenase) in the superficial gastrocnemius of PGC-1β^(i)skm-/-^ mice (two-way ANOVA genotype main effect p<0.0001, F=73.64) (Figure 3A). Moreover, treatment with atorvastatin was associated with a decrease of NADH dehydrogenase staining of the gastrocnemius muscle of WT and KO mice compared to their respective control group (two-way ANOVA treatment main effect p<0.0001, F=166.13), leading to additive effects of these two parameters (two-way ANOVA interaction effect p=0.0027, F=13.66) (Figure 3A). The percentage of stained fibers was also decreased in the oxidative vastus intermedius muscle of KO-ATO mice compared to KO-CTL, but not in the WT-ATO mice (two-way ANOVA interaction effect p=0.0033, F=14.72) (Figure 3B). We observed no difference in NADH dehydrogenase staining between control groups in the oxidative muscle.

**Figure 3:**
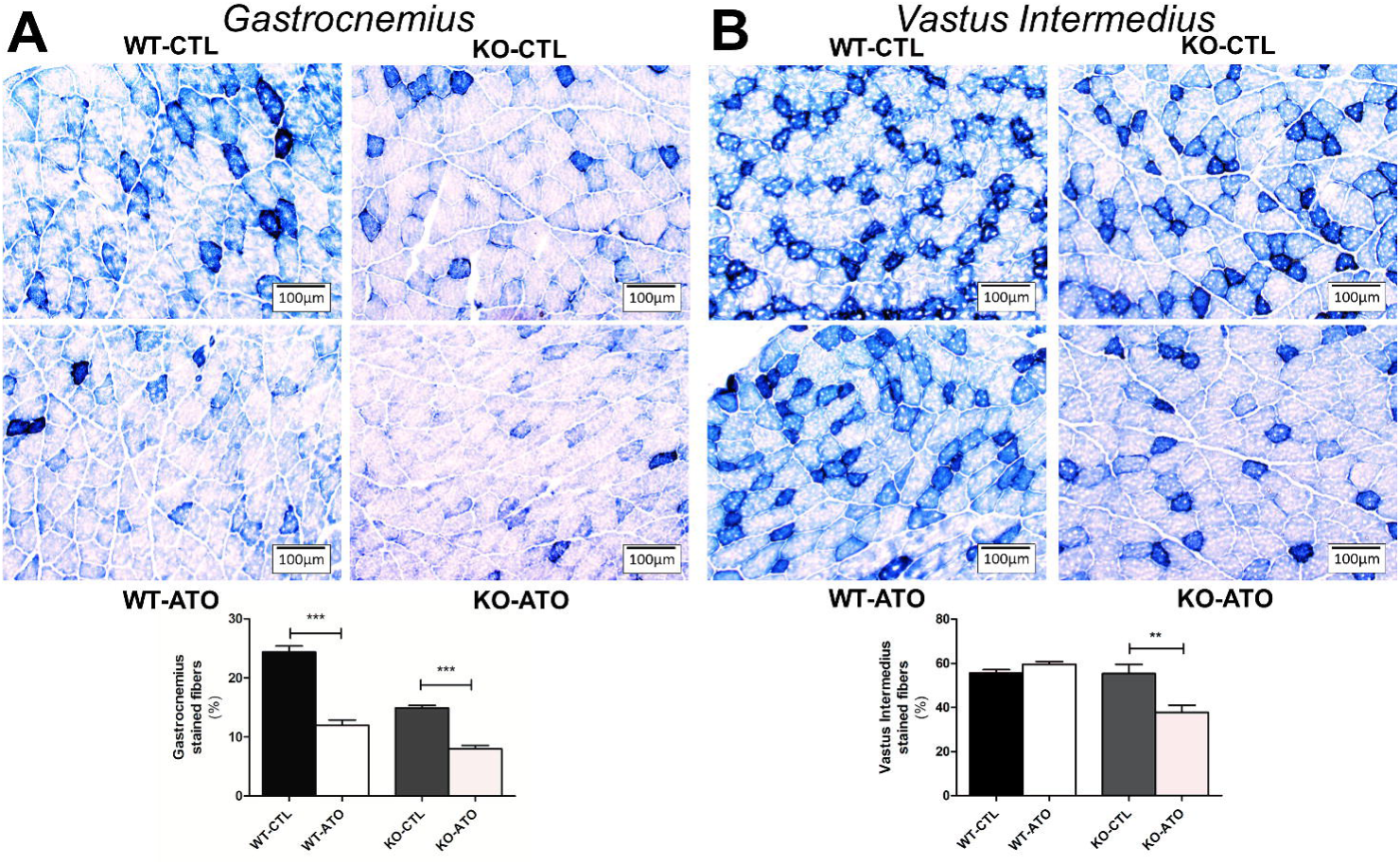
Atorvastatin impaired muscle phenotype in fiber type in the glycolytic skeletal muscle and in the oxidative skeletal muscle of PGC-1β deficient mice. **(A-B)** NADH diaphorase staining, and stained fibers percentage of **(A)** superficial gastrocnemius muscle sections, and **(B)** vastus intermedius muscle sections. Results are expressed as mean±SEM; n=8−13 per group, **p<0.01, ***p<0.001.

Based on these results, we determined the mitochondrial respiration rates in permeabilized fibers from both muscle types. Atorvastatin treatment for two weeks impaired the glycolytic muscle mitochondrial function in both WT and KO mice (Figure 4A). Although no differences were found in the basal respiratory rates, atorvastatin treatment decreased OXPHOS C_I_-linked substrate state O_2_ consumption in both WT-ATO and KO-ATO groups compared to their respective control group (two-way ANOVA treatment main effect p=0.0008, F=13.82). In addition, maximal OXPHOS respiratory rates (C_I+II_-linked substrate state) (two-way ANOVA treatment main effect p=0.0011, F=7.19) and C_II_-linked substrate state (two-way ANOVA treatment main effect p=0.0413, F=4.45) were decreased by atorvastatin in PGC-1β^(i)skm-/-^, but not in the respective control mice. In comparison, no significant differences were found in the Respiratory Control Ratio (RCR) of the gastrocnemius (Figure 4B). In the oxidative soleus muscle, mitochondrial function was impaired by atorvastatin only in KO mice (Figure 4C). Basal mitochondrial respiration as well as C_I_-linked substrate state (two-way ANOVA interaction effect p=0.0459, F=2.26), C_I+II_-linked substrate state (two-way ANOVA interaction effect p=0.0436, F=4.35), and C_II_-linked substrate state (two-way ANOVA interaction effect p=0.0307, F=5.03) were decreased in soleus of KO-ATO compared to KO-CTL mice. Again, no differences were found in the soleus RCR (Figure 4D).

**Figure 4:**
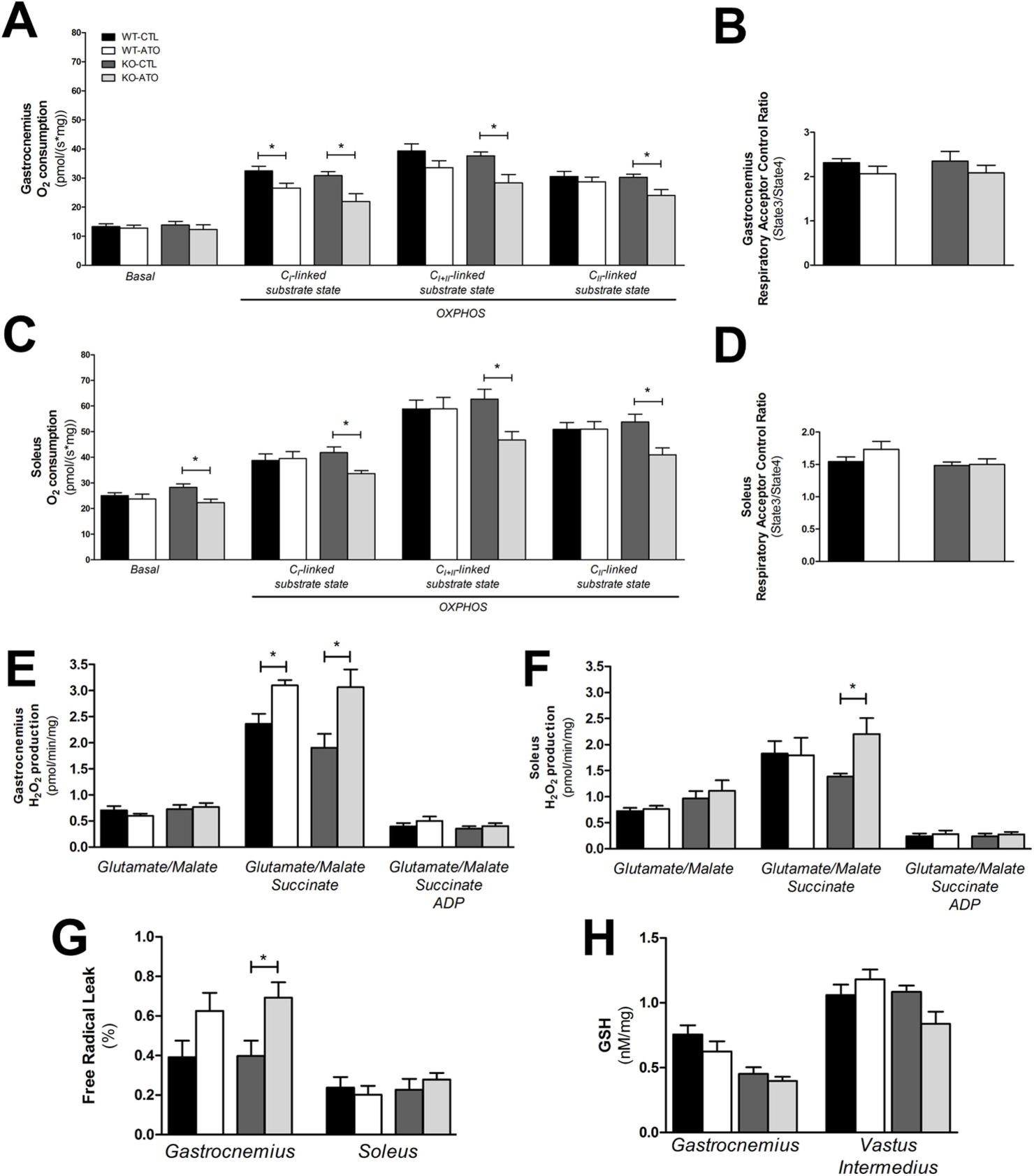
Impaired mitochondrial respiratory chain and enhanced ROS production. Mitochondrial oxidative capacities **(A&C)**, and mitochondrial respiratory ratios **(B&D)** of **(A-B)** superficial gastrocnemius muscle, and **(C-D)** soleus muscle. **(E-F)** Mitochondrial H_2_O_2_ production of **(E)** superficial gastrocnemius muscle, and **(F)** soleus muscle. **(G)** Mitochondrial free radical leak. **(H)** Muscle content of reduced glutathione (GSH). Results are expressed as mean ±SEM; n=8−13 per group, *p<0.05.

If the above-mentioned hypothesis were correct, the observed impairment in mitochondrial function should be associated with increased mitochondrial H_2_O_2_ production, especially under reverse electron flux conditions. Mitochondrial H_2_O_2_ production was increased in the glycolytic gastrocnemius muscle, in both WT-ATO and KO-ATO groups (two-way ANOVA treatment main effect p=0.0006, F=15.17) compared to their respective control group under reverse electron flux conditions (Figure 4E). In the oxidative soleus, H_2_O_2_ production was also increased in KO-ATO compared to KO-CTL mice (Figure 4F). These measurements enabled us to calculate the Free Radical Leak (FRL), which was increased in the atorvastatin-treated groups (two-way ANOVA treatment main effect p=0.0016, F=11.88, DF=1), especially in the KO-ATO group compared to KO-CTL in the glycolytic muscle (Figure 4G). Although the cellular reduced glutathione (GSH) pool, an important constituent of the antioxidative defense system, was affected in the gastrocnemius in the PGC-1β^(i)skm-/-^ mice (two-way ANOVA genotype main effect p=0.0003, F=19.93), it only tended to decrease in WT-ATO and KO-ATO mice compared to their respective control. In vastus intermedius, atorvastatin treatment affected the GSH pool differently in function of the mouse genotype (two-way ANOVA interaction effect p=0.0376, F=5.09), with deleterious effects in the KOATO group (Figure 4H).

### Mitochondrial adaptive pathways were involved for statin tolerance

To determine the consequences of oxidative stress, we then investigated the mitochondrial biogenesis pathway in more detail. In the glycolytic gastrocnemius muscle, we observed an impairment of the mitochondrial biogenesis pathway in WT-ATO compared to WT-CTL mice with a significant decrease of PGC-1α and PGC-1β mRNA levels, whereas the decrease in the mRNA levels of NRF1, TFAm and NRF2 was not statistically significant (Figure 5A). A similar impairment was detected in KO-ATO compared to KO-CTL mice, with a decrease of PGC-1β (which detected levels reflect its expression in satellite cells), NRF2, and TFAm mRNA expression, whereas the decrease in PGC-1α and NRF1 mRNA expression was not statistically significant. In addition, the two-way ANOVAs revealed a main effect of atorvastatin in both genotypes, with a repression of the whole mitochondrial biogenesis pathway: PGC-1α (p=0.0114, F=8.67), PGC-1β (p=0.0041, F=12.50), NRF1 (p=0.0104, F=8.76), NRF2 (p=0.0041, F=11.74), and TFAm (p=0.0033, F=12.20). In contrast, in the oxidative vastus intermedius muscle (Figure 5B), we observed an activation of mitochondrial biogenesis pathways, with an increased expression of PGC-1α, NRF1, NRF2, and TFAm in WT-ATO compared to WT-CTL mice. On the contrary, in the KO-ATO group, we observed a repression of the mitochondrial biogenesis pathway in the oxidative vastus intermedius muscle, with reduced mRNA expression levels of PGC-1α, PGC-1β, and TFAm compared to KO-CTL mice. As a consequence, the two-way ANOVAs revealed an interaction effect for PGC-1α (p=0.0062, F= 9.92), NRF1 (p=0.0078, F=9.11), NRF2 (p=0.0241, F=6.07), and TFAm (p=0.0016, F=14.31).

**Figure 5:**
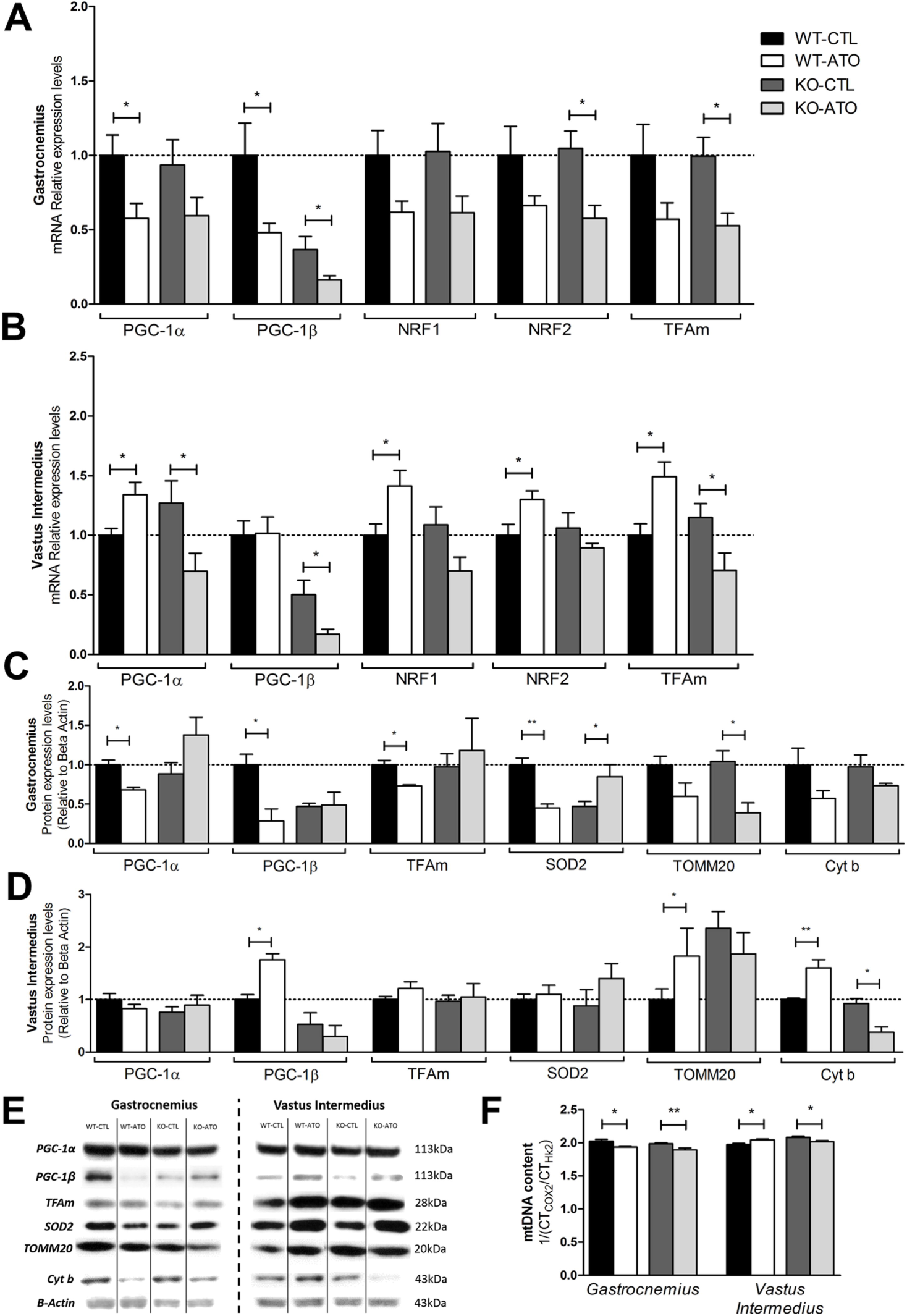
Different effects of atorvastatin on mitochondrial biogenesis. **(A)** mRNA transcript levels in superficial gastrocnemius **(B)** and in vastus intermedius muscle. Values are relative to 18S expression. **(C)** Muscle protein expression levels in superficial gastrocnemius **(D)** and vastus intermedius muscle. Values are relative to β-actin expression. **(E)** Immunoblots of PGC-1β, TFAm, TOMM20, cytochrome b and β-actin. **(F)** Mitochondrial DNA quantification. Results are expressed as mean ±SEM; n=8−13 per group, *p<0.05, **p<0.01.

The study of mitochondrial adaptations by western blotting grossly confirmed what we had shown at the mRNA level. In the glycolytic gastrocnemius muscle (Figure 5C & E), we observed in WT-ATO mice a decrease of mitochondrial biogenesis-linked proteins such as PGC-1α, PGC-1β and TFAm compared to WT-CTL mice. As previously described, SOD2 expression was decreased in the glycolytic muscle of KO compared to WT mice (31). This repression of mitochondrial biogenesis was supported by a trend to decrease in the expression of TOMM20, and of cytochrome b (Figure 5C) and a significant decrease in the mtDNA content in WT-ATO compared to WT-CTL mice (Figure 5F). In the gastrocnemius muscle of KO-ATO mice, although we did not observe a decrease in mitochondrial biogenesis-linked proteins, we noticed a decrease in TOMM20 and a trend for a decrease in cytochrome b expression (Figure 5C) as well as a decrease in the mtDNA content compared to KO-CTL mice (Figure 5F). This was confirmed by the two-way ANOVA results, which revealed a main effect of atorvastatin for TOMM20 (p=0.0030, F=15.21), cytochrome b (p=0.0329, F=6.34), and mtDNA (p=0.0003, F=19.74). In the oxidative vastus intermedius muscle, we observed in WT-ATO an activation of the mitochondrial biogenesis pathway with an increase in PGC-1β compared to WT-CTL mice (Figure 5D). This was confirmed with the increase in TOMM20 and cytochrome b expression and in the mtDNA content (Figure 5F). In KO-ATO mice, although we did not observe a decrease in mitochondrial biogenesis-linked proteins, we observed a decrease in cytochrome b protein expression (Figure 5D), and in the mtDNA content compared to KO-CTL mice (Figure 5F). These observations were confirmed by the two-way ANOVA results, which revealed an interaction effect for cytochrome b (p=0.0002, F=34.70), and mtDNA (p=0.0008, F= 15.40).

### PGC-1β protected the oxidative muscle from statin-induced apoptosis

Western blot analysis of caspase 3 (Figure 6A) revealed in the glycolytic gastrocnemius an effect of atorvastatin treatment (two-way ANOVA treatment main effect p=0.0001, F=21.01), with a 2-fold increase in the cleaved caspase 3/total caspase 3 ratio in WT-ATO compared to WT-CTL, and a 2.2-fold increase for KO-ATO compared to KO-CTL. In the oxidative muscle, no differences were observed between WT-CTL and WT-ATO, but this ratio was increased by 2.2-fold for KO-ATO compared to KO-CTL mice (two-way ANOVA interaction effect p=0.0173, F=6.73). To confirm the induction of apoptotic pathways by atorvastatin, we performed a TUNEL staining (Figure 6B-D), which revealed a similar effect in the glycolytic skeletal muscle (two-way ANOVA treatment main effect p=0.0002, F=38.69), and in the oxidative skeletal muscle (two-way ANOVA interaction effect p=0.0005, F=32.54). In WT mice, TUNEL staining data showed, as reported previously in rats (13), that atorvastatin increased the proportion of myonuclei positive for DNA strand breaks in the glycolytic (Figure 6B & D) but not in the oxidative skeletal muscle (Figure 6C & D). In contrast to these observations, in KO-ATO mice, both glycolytic and oxidative-type skeletal muscles displayed a higher percentage of apoptotic nuclei.

**Figure 6:**
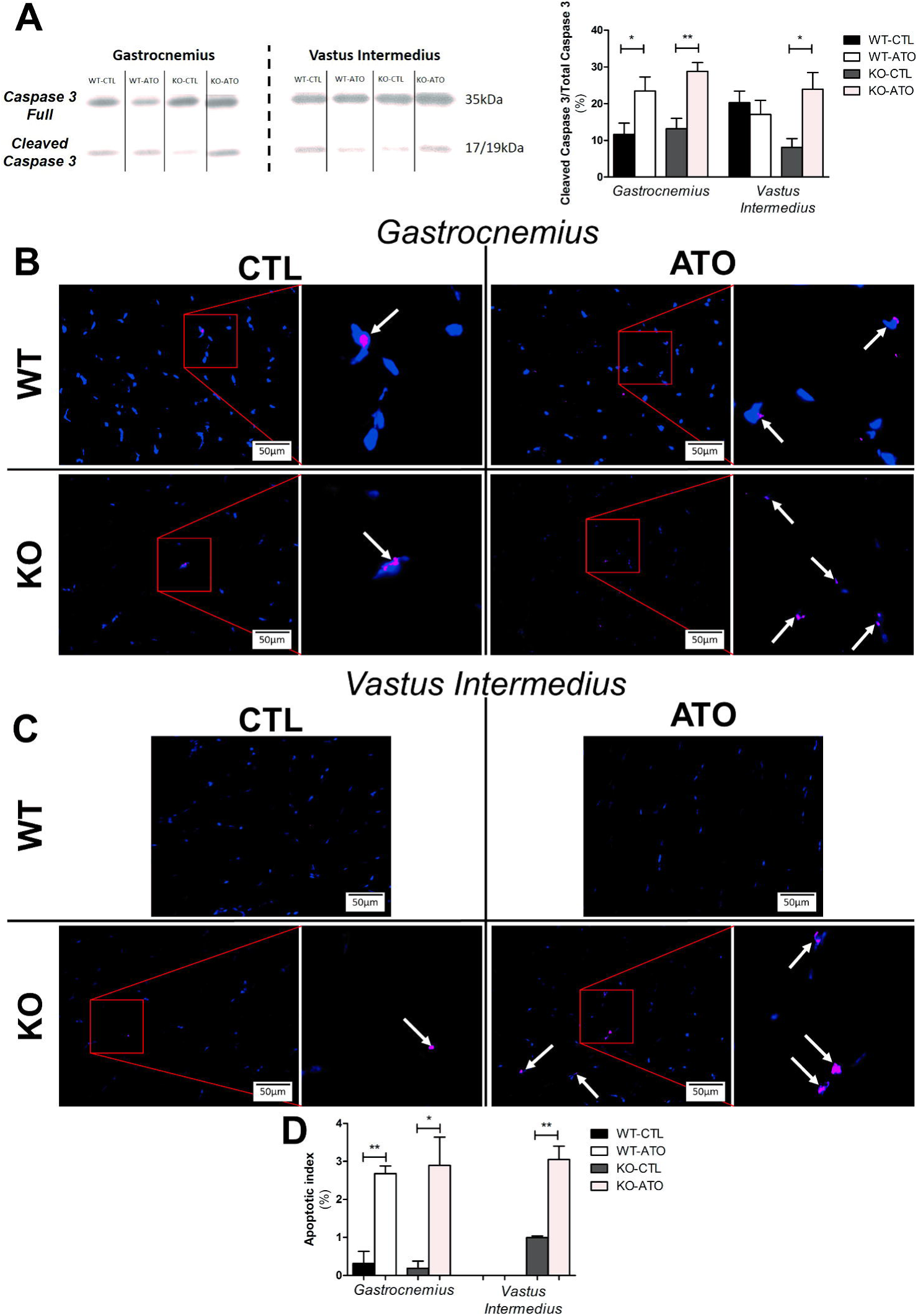
Treatment with atorvastatin induces apoptotic pathways. **(A)** Immunoblots of caspase 3 and cleaved caspase 3, and cleaved caspase 3 expression relative to caspase 3 expression. **(B-C)** Nuclei were stained with DAPI (blue), and TUNEL nuclei were visualized with magenta fluorescein. Merged DAPI and TUNEL nuclei from different groups are shown and white arrows indicating apoptotic nuclei. Adjacent pictures correspond to a magnification of the red square area. Merged TUNEL images of **(B)** superficial gastrocnemius and **(C)** vastus intermedius muscle sections. **(D)** Apoptotic index calculated from TUNEL staining. Results are expressed as mean ±SEM; n=8−13 per group, *p<0.05, **p<0.01.

## Discussion

This study provides four major findings: 1) Atorvastatin treatment induced a switch in fiber type in the glycolytic skeletal muscle and in the oxidative skeletal muscle of PGC-1β deficient mice, 2) Atorvastatin treatment exhibited mitochondrial dysfunction and apoptosis in the glycolytic skeletal muscle and in the oxidative skeletal muscle of PGC-1β deficient mice, 3) Atorvastatin exhibited a beneficial effect by triggering mitochondrial biogenesis in wild-type oxidative muscles, and 4) The loss of PGC-1β rendered oxidative muscle as sensitive as glycolytic muscle towards statin-associated myotoxicity (Figure 7). These findings provide a new mechanism of how statins affect the integrity of skeletal muscle according the muscular phenotype and demonstrate the importance of the presence of PGC-1β for the defense of oxidative muscle against the myotoxic effects of statins.

**Figure 7:**
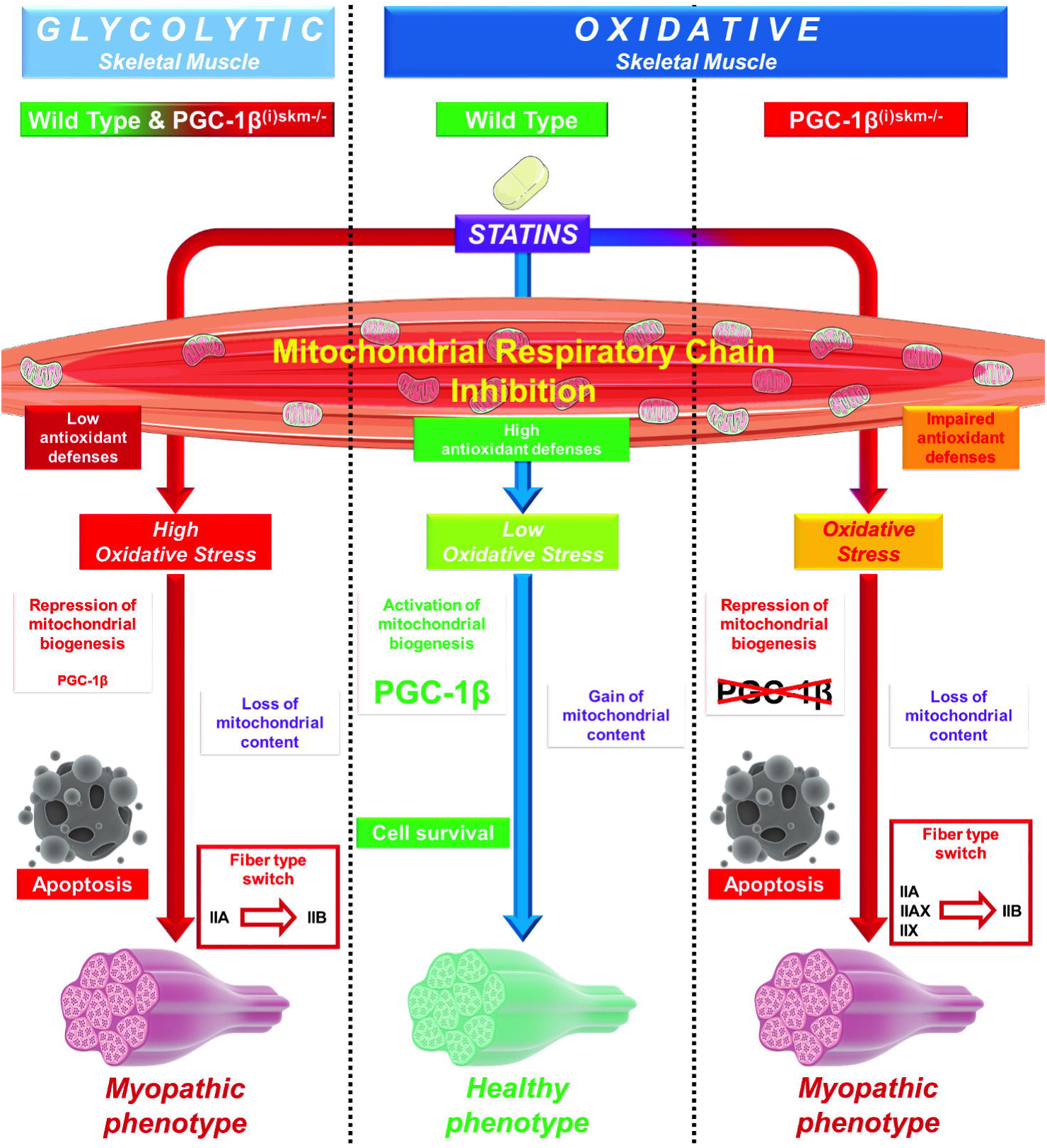
Proposed scheme illustrating the action of statins on mitochondrial function in wild type and PGC-1β deficient mice according to muscular phenotype. Statins impair mitochondrial function in the glycolytic skeletal muscle from wild-type and PGC-1β^(i)skm-/-^ mice, leading to the formation of oxidative stress, the triggering of apoptotic pathways, and a shifting towards highly glycolytic fiber type. Oxidative muscle contains higher antioxidant defences that protect this muscle from the action of statins, leading to cell survival. However, the loss of PGC-1β renders the oxidative muscle susceptible to statins leading to the appearance of oxidative stress and subsequently to apoptosis.

We first showed that atorvastatin treatment decreased the grip strength in wild-type and PGC-1β^(i)skm-/-^ mice. This decrease was correlated with an increase of creatine kinase in the ATO-treated mice compared to CTL mice. Although the absence of PGC-1β in PGC-1β^(i)skm-/-^ mice exacerbated atorvastatin-induced CK release, it was not associated with an increase in the reduction of the muscle peak force, suggesting that atorvastatin itself could cause this toxicity. Our data showed a decreased muscle fiber area in glycolytic muscle from both mice models. These results in the wild-type groups are in line with what has been previously described by us and by others (8,35), showing statin-induced muscular atrophy in glycolytic muscle. Hanai et al. described an increased atrogin-1 mRNA expression in patients with statin-induced muscle injury (8). A similar increase was also observed in C2C12 myoblasts at the mRNA and at the protein level (35). In addition, Bonifacio et al. showed a decrease in the phosphorylation of FoxO3, which is responsible for the increase in atrogin-1 expression and for the induction of the atrophic phenotype. Interestingly we also detected a decrease of muscle fiber area in oxidative muscle of PGC-1β^(i)skm-/-^ mice. These results could possibly be explained by the fact that PGC-1β has been reported to suppress protein degradation by decreasing FoxO3 transcriptional activity (36), and that reduced PGC-1β expression is associated with muscle wasting (37).

Contrary to glycolytic skeletal muscle, oxidative skeletal muscle appears to be protected from the harmful effects of statins (13,38,39). Oxidative muscle is mainly composed of type I slow-twitch and type IIA fast-twitch fibers, whereas glycolytic muscle contains mainly type IIB and type IIX slow-twitch fibers (40). Although overexpression of PGC-1β has been shown to promote the formation of type IIX fibers in skeletal muscle (41), PGC-1β deficiency did not affect fiber type composition (31). These data showed that the PGC-1β deficient oxidative vastus intermedius muscle, when exposed to atorvastatin, developed a similar myofiber phenotype as the glycolytic gastrocnemius muscle. To date, the molecular regulation of fiber-type determination is not completely understood. Factors proposed to drive the formation of oxidative myofibers include AMP-activated protein kinase (AMPK) and PGC-1α (42,43) but not PGC-1β (31). On the other hand, glycolytic muscle determination has been described to result from activation of Akt through a mechanism involving Baf60c, a transcriptional cofactor enriched in fast-twitch muscle, and DEP domain-containing mTOR-interacting protein (44). In our previous study we showed however, that the activation of Akt is impaired by statins in C2C12 cells mainly due to a reduced mTORC2 activity (9). The observed shift from type IIA to type IIB fibers in PGC-1β^(i)skm-/-^ mice treated with atorvastatin may therefore involve different mechanisms than activation of Akt.

We observed an impaired mitochondrial function in glycolytic and oxidative muscles of KO mice, but only in glycolytic muscle in WT. Mitochondria appear to adapt better to the toxic stress associated with statins in skeletal muscle with an oxidative phenotype than with a glycolytic one. Maximal OXPHOS respiratory rate depends on convergent electron flow through complexes I & II to the Q-junction of the electron transport chain (45). Biosynthesis of CoQ_10_ is dependent on mevalonate production within the cell. Statins potentially decrease the biosynthesis of ubiquinone by inhibiting mevalonate production. This decrease in ubiquinone formation could therefore be a plausible explanation for the decrease in mitochondrial oxygen consumption. In the current study, the serum atorvastatin concentrations were not different between WT-ATO and KO-ATO mice, suggesting also similar concentrations in skeletal muscle and similar effects on ubiquinone biosynthesis. Since we observed impaired function of the electron transport chain only in skeletal muscle with a glycolytic, but not with an oxidative phenotype, we considered a decrease in the skeletal muscle ubiquinone content an unlikely explanation for mitochondrial dysfunction. In support of this assumption, in rats with muscle damage due to treatment with increasing doses of cerivastatin, the skeletal muscle ubiquinone content was not different from vehicle-treated control rats (46). In addition, the data regarding the effect of CoQ_10_ supplementation in patients with statin-associated myopathy are inconsistent (47). It appears that the skeletal muscle CoQ_10_ content remains high enough in most patients treated with statins to maintain the electron transport chain function. Interestingly, it has recently been reported that statin-induced myopathy is associated with a direct inhibition of mitochondrial complex III at the Q_0_ ubiquinone site level (11), which is known to be a site of mitochondrial ROS generation (48). In addition, if the Q_0_ ubiquinone binding site at complex III is impaired, complex II delivers a higher quantity of electrons to ubiquinone than complex III can accept, leading to an “over-reduction” of ubiquinone. This favours a reverse electron flux towards complex I (49), putting complex I in a reduced state and promoting the production of ROS.

Hence, both complex I and III are ROS production sites potentially involved in statin myotoxicity, possibly further damaging the enzyme complexes of the mitochondrial respiratory chain. These results are in line with data published by us and others (10,13). Indeed, Kwak et al. observed an increase in mitochondrial ROS production in primary human myotubes treated with simvastatin due to inhibition of the mitochondrial respiratory chain. In support of the current findings, in L_6_ myoblasts treated with atorvastatin, we recently demonstrated simultaneous inhibition of the mitochondrial respiratory chain and mitochondrial ROS production in a dose-dependent manner (13). In addition, we also observed an increased ROS production in both human deltoid biopsies and in the glycolytic skeletal muscle in rats. Our results show that in WT mice a mild increase in ROS levels by atorvastatin induced the expression of PGC-1β in oxidative muscle. However, when PGC-1β expression is absent in the same muscles, the statin-associated ROS production is higher, leading to a down-regulation of mitochondrial biogenesis, and to the triggering of apoptotic pathways. It has previously been demonstrated that exposure of cells to H_2_O_2_ increases PGC-1β expression in cells (50), and we have shown that PGC-1β protects C2C12 cells from H_2_O_2_-induced cell death (31).

Inhibition of mitochondrial biogenesis by statins has also been demonstrated in patients, since skeletal muscle mitochondrial DNA was decreased in patients with statin-induced myopathy (16). Our results in glycolytic muscle of WT and KO mice and in oxidative muscle of KO mice are in line with these observations. On the other hand, we observed surprisingly a triggering of the mitochondrial biogenesis pathway in oxidative skeletal muscle of WT mice treated with atorvastatin, leading to an increased mitochondrial content, which was possibly explained by mitochondrial hormesis. Our study demonstrates the important role of PGC-1β in triggering antioxidative defense and mitochondrial proliferation in oxidative muscle in the presence of a mitochondrial toxicant such as atorvastatin. We previously showed that statins had opposite effects on mitochondria of cardiac and skeletal muscle (22). Indeed, we observed that statins had deleterious effects on skeletal muscle and beneficial effects on the cardiac muscle by triggering the mitochondrial hormesis phenomenon. In a more recent publication, we showed that statins can trigger mitochondrial apoptotic pathways in glycolytic skeletal muscle, whereas oxidative skeletal muscle was resistant (13). Moreover, we recently showed *in vitro* that increasing the mitochondrial content by triggering mitochondrial hormesis could protect L_6_ myoblasts from the deleterious effects of statins (14).

In the present study, we confirm that the mitochondrial content is an important factor in preventing statin-associated toxicity. Indeed, statins seem to have beneficial effects not only in the heart, but also in skeletal muscles displaying an oxidative metabolism with a high mitochondrial content. On the other hand, in glycolytic muscles which possess a lower mitochondrial content, statins trigger apoptotic pathways in a mitochondrial ROS-dependent manner. In addition, we show the importance of the ability to adapt the mitochondrial biogenesis pathways to counteract statin-associated toxicity in a long-term setting. Mitochondrial function and adaptation capacities could hence partially explain why some patients are developing a statin-induced myopathy, and some are tolerant to statins. These data underscore the importance of mitochondrial biogenesis for statin tolerance, and define a hinge role of PGC-1β in mitochondrial adaptation to oxidative stress.

## Methods

### Animals

Experiments were performed on adult mice (18-23 weeks at the beginning of the study; ~30-35g). Generation of PGC-1β^(i)skm-/-^ mice, in which PGC-1β is selectively ablated in skeletal muscle myofibers at adulthood, has been described previously (31). Briefly, PGC-1β^(i)skm-/-^ were bred with C57BL/6 HAS-Cre-ER^T2(tf/0)^ transgenic mice that express the tamoxifen-dependent Cre-ER^T2^ recombinase under the control of the HAS regulatory elements, and their offsprings were intercrossed to generate HSA-Cre-ER^T2(tg/0)^/PGC-1β^L2/L2^ pre-mutant mice and HSA-Cre-ER^T2(0/0)^/PGC-1^βL2/L2^ control mice. Pre-mutant male mice and sex-matched control littermates were intraperitoneally injected with tamoxifen (1 mg per day, for 5 days) at 7 weeks of age.

Animals were housed in neutral temperature environment (22° ± 2 °C) on a 12:12 hour photoperiod and had free access to food and water. Breeding and maintenance of mice were performed in the accredited IGBMC/ICS animal house (C67-2018-37 notification of 16/10/2013), in compliance with French and EU regulations on the use of laboratory animals for research, under the supervision of D. M. who holds animal experimentation authorizations from the French Ministry of agriculture and Fisheries (N°67-209 and A 67-227). All animal experiments were approved by the Ethics Committee Com’Eth (Comité d’Ethique pour l’Expérimentation Animale, Strasbourg, France). Atorvastatin (Tahor^®^) was generously provided by Pfizer. 43 mice were randomly divided in 4 groups as follows: (1) control animals (WT-CTL, n=13); (2) animals treated with atorvastatin 5 mg/kg/day in drinking water for two weeks (WT-ATO, n=11); (3) PGC-1β^(i)skm-/-^ mice (KO-CTL, n=8); (4) PGC-1β^(i)skm-/-^ treated with atorvastatin 5 mg/kg/day in drinking water for two weeks (KO-ATO, n=11).

### Sample collection

Animals were killed by cervical dislocation and tissues were immediately collected, weighed, and a part of each muscle tissue was immediately frozen in isopentane cooled by liquid nitrogen and stored at −80°C for later analysis, or processed for biochemical and histological analysis. The oxidative soleus muscle and the glycolytic superficial part of the gastrocnemius muscle were excised and cleaned of adipose and connective tissues. We conducted our experiments on different muscles, separating them by muscle type. Muscles are classified by mitochondrial mass: we used the superficial part of the gastrocnemius muscle, which is mostly glycolytic, with a low mitochondrial content. For the oxidative-type muscles with a high mitochondrial content, we studied the soleus muscle, and the vastus intermedius: the most oxidative part of the quadriceps, which is located close to the femur bone.

### Plasma biochemistry

Blood was collected by cardiac puncture immediately after death in ethylenediaminetetracetic acid (EDTA)-rinsed tubes and then centrifuged at 1100 g for 15 min at 4°C. The plasma was separated and stored at −80°C until analysis. The analysis of plasma biochemical parameters was performed on randomly selected plasma samples. Plasmatic creatine kinase activity was determined using the Creatine Kinase Activity Assay Kit from Sigma-Aldrich (MAK116; St Louis, USA), according to the manufacturer’s instructions.

### Atorvastatin plasma concentrations

Atorvastatin was analyzed on an UHPLC system (Shimadzu, Kyoto, Japan) connected to an API 5500 tandem mass spectrometer (AB Sciex, Ontario, Canada). The UHPLC system consisted of four LC-30AD pumps, a SIL-30AC autosampler, a CTO-20AC column oven, a DGU-20A5 degassing unit, and a CBM-20A controller. Chromatography was performed on a kinetex 2.6 µ F5 100 Å (50x2.1 mm) analytical column (Phenomenex, Torrance, USA). Mobile phase A consisted of water plus 0.1% formic acid, while mobile phase B was methanol supplemented with 0.1% formic acid. The following gradient program of mobile phase B was applied: 40% (0-0.25 min), 95% (0.25-1.5 min), 95% (1.5-2.0 min), 40% (2.0-2.25 min). The flow rate was set at 0.4 mL/min at 40°C. The introduced sample was pre-column diluted with water 0.1% formic acid during the first 0.25 min of each run. Atorvastatin eluted after 1.7 minutes, hence the UHPLC was only connected from minute 1-2 with the mass spectrometer to avoid superfluous contamination of the system. Atorvastatin and atorvastatin-d5 were detected by multiple reaction monitoring using electrospray ionization in the positive mode. A mass transition of 559.1→440.1 m/z and 564.1→445.1 was used for atorvastatin and atorvastatin-d5, respectively (dwell time: 25 msec, declustering potential: 126 V, entrance potential: 10 V, collision energy: 33 V, cell exit potential: 30 V). The mass spectrometer was operated at an ion source gas 1 of 45 l/min (N2), an Ion source gas 2 of 60 l/min (N2), a curtain gas of 30 l/min, a collision gas set to medium (N2), an ion spray voltage of 4500 V, and a source temperature of 350°C. Analyst software 1.6.2 (AB Sciex, Ontario, Canada) was used to control the LC-MS/MS system. Calibration lines were prepared in blank mouse plasma. Thereby, blank plasma was spiked with atorvastatin at a final concentration of 2242 nM and serially diluted to 4.4 nM (≤1% DMSO). Plasma aliquots of 10 µL were precipitated with 150 µL methanol 0.1% formic acid containing the internal standard (atorvastatin-d5: 25 nM). Samples were mixed with a VX-2500 multi-tube vortexer (VWR, Dietikon, Switzerland) for about 1 min and centrifuged at 4000 rpm during 30 min (Eppendorf 5810R, Hamburg, Germany). 10 µL supernatant were injected into the LC-MS/MS system.

### Grip strength Test

A Grip Strength Meter (Bioseb, Vitrolles, France) was used to measure combined forelimb and hindlimb grip strength. The test was repeated three consecutive times within the same session, and the mean value was recorded as the maximal grip strength for each mouse. All tests were performed by the same operator.

### Histology

The analysis of all histological parameters was performed on randomly selected slides samples. Hematoxylin & Eosin staining was performed on 10 µm frozen sections of 3-4 samples per group on the glycolytic muscle gastrocnemius, and the more oxidative type vastus intermedius muscle. Hematoxylin & Eosin staining photographs were captured on an Olympus IX83 microscope (Olympus, Hamburg, Germany). Evaluation and interpretation of the results were realized according to the MDC1A_M.1.2.004 SOP. Fiber area was determined using the cellSens Dimension software (Olympus, Hamburg, Germany) on at least 15 fibers per picture, and the mean of three independent pictures was calculated for each sample. Measurements were made on pictures obtained with a 20X objective. Illustrations provided in the figures have been obtained with a 40X objective. Presence of central nuclei was confirmed by color deconvolution using the ImageJ software (National Institute of Health) in order to exclude staining artifacts.

NADH diaphorase staining was performed on 10 µm frozen sections of 3-4 samples per group, on the glycolytic muscle gastrocnemius, and the more oxidative type vastus intermedius muscle. NADH staining photographs were captured on an Olympus IX83 microscope (Olympus, Hamburg, Germany, 10X and 20X objectives). Determination of the percentage of oxidative-type fibers was performed by manual counting on pictures obtained with a 10X objective. Illustrations provided in the figures have been obtained with a 20X objective.

### Myosin heavy chain expression immunofluorescence

Immunofluorescence analysis of MHC expression was performed as described by (51), on 10 µm frozen sections of 3 samples per group on the glycolytic muscle gastrocnemius, and the more oxidative type vastus intermedius muscle. Primary antibodies against MHCI (BA-F8, 1/50), MHCIIA (SC-71, 1/600), and MHCIIB (BF-F3, 1/100) were purchased from the Developmental Studies Hybridoma Bank (University of Iowa), whereas secondary antibodies were purchased from ThermoFisher scientific (A21140: Goat anti-mouse IgG2b secondary antibody, Alexa Fluor^®^ 350 conjugate; A21121: Goat anti-mouse IgG1 secondary antibody, Alexa Fluor^®^ 488 conjugate; and A21426: Goat anti-mouse IgM heavy chain secondary antibody, Alexa Fluor^®^ 555 conjugate). All antibody cocktails were prepared in block solution (10% goat serum (Gibco) in PBS). Briefly, slides were incubated with block solution for one hour. Primary antibody cocktail was then applied, and slides were incubated for two hours. After PBS wash, secondary antibody cocktail was applied, and slides were incubated for one hours. After PBS wash, slides were mounted with coverslips, using Prolong^®^ Gold antifade reagent (ThermoFisher scientific, Waltham, MA, USA). MHC immunofluorescence pictures were captured on an Olympus IX83 microscope (Olympus, Hamburg, Germany). Determination of the percentage of type I, IIA, IIAX, IIX, IIXB, and IIB fibers was performed by manual counting on pictures obtained with a 20X objective. Merged illustrations provided in the figures have been obtained with a 40X objective.

### Study of muscle mitochondrial respiration

This technique ensured determination of global mitochondrial function, reflecting both the density as well as the functional properties of the muscle mitochondria (52). The mitochondrial respiration was studied from saponin-skinned fibers that keeping mitochondria in their architectural environment, in superficial gastrocnemius, and soleus muscles. The analysis took place in a thermostated oxygraphic chamber at 37°C with continuous stirring (Oxygraph-2k, Oroboros instruments, Innsbruck, Austria). Approximately 2 mg of fibers were placed in respiration medium (2.77 mM CaK_2_EGTA, 7.23 mM K_2_EGTA, 6.56 mM MgCl_2_, 20 mM imidazole, 20 mM taurine, 0.5 mM dithiothreitol, 50 mM K-methane sulfonate, 5 mM glutamate, 2 mM malate, 3 mM phosphate, and 2 mg/mL of Bovine Serum Albumin [BSA]; pH=7) in the oxygraphic chamber. After the determination of the basal oxygen consumption with glutamate (5 mM) and malate (2 mM) (Basal), OXPHOS C_I_-linked substrate state was measured in the presence of saturating amount of adenosine diphosphate (2 mM ADP). When OXPHOS C_I_-linked substrate state was recorded, electron flow went through complexes I, III, and IV The maximal OXPHOS respiration rate C_I+II_-linked substrate state was then measured by adding succinate (25 mM). Complex I was blocked with rotenone (0.5 µM), allowing to measure OXPHOS C_II_-linked substrate state. Respiratory rates were expressed as pmol O_2_ x s^−1^ x mg^−1^ wet weight. Respiratory Acceptor Control Ratio (RCR) was determined by calculating State 3/State 4 respiratory rates.

### Mitochondrial H_2_O_2_ production in permeabilized fibers

H_2_O_2_ production was studied from saponin-skinned fibers that keep mitochondria in their architectural environment, in superficial gastrocnemius, and soleus muscles. The permeabilized bundles were placed in ice-cold buffer Z containing 110 mM K-methane sulfonate, 35 mM KCl, 1 mM EGTA, 5 mM K_2_HPO_4_, 3 mM MgCl_2_, 6 mM H_2_O, 0.05 mM glutamate, and 0.02 mM malate with 0.5 mg/ml BSA (pH 7.1, 295 mOsmol/kg H_2_O). H_2_O_2_ production was measured with Amplex Red (Invitrogen Life Technologies, Rockville, MD, USA), which reacted with H_2_O_2_ in a 1:1 stoichiometry catalyzed by HRP (horseradish peroxidase; Invitrogen Life Technologies, Rockville, MD, USA) to yield the fluorescent compound resorufin and a molar equivalent of O_2_ (53). Resorufin has excitation and emission wavelengths of 563 nm and 587 nm, respectively, and is extremely stable once formed. Fluorescence was measured continuously with a Fluoromax 3 (Jobin Yvon) spectrofluorometer with temperature control and magnetic stirring. After a baseline (reactants only) was established, the reaction was initiated by adding a permeabilized fiber bundle to 600 µL of buffer Z. Buffer Z contained 5 mM Amplex Red, 0.5 U/mL HRP, 5 mM glutamate, and 2 mM malate as substrates at 37°C. Succinate (25 mM) was then added for the measurement of H_2_O_2_ production under reverse electron flux condition. ADP (2mM) was added in order to determine C_I+II_-linked substrate state H_2_O_2_ production for the determination of the Free Radical Leak (FRL) At the conclusion of each experiment. The results were reported in pmol H_2_O_2_ x s^−1^ x mg^−1^ wet weight.

### Free Radical Leak

H_2_O_2_ production and O_2_ consumption were measured in parallel under similar experimental conditions (C_I+II_-linked substrate state). This allowed the calculation of the fraction of electrons out of sequence which reduce O_2_ to ROS in the respiratory chain (the percentage of free radical leak) instead of reaching cytochrome oxidase to reduce O_2_ to water (53). Because two electrons are needed to reduce one mole of O_2_ to H_2_O_2_, whereas four electrons are transferred in the reduction of one mole of O_2_ to water, the percent of FRL was calculated as the rate of H_2_O_2_ production divided by twice the rate of O_2_ consumption, and the result was multiplied by 100.

### Quantitative Real Time Polymerase Chain Reaction (qRT-PCR)

Total RNA was obtained from superficial gastrocnemius and the oxidative part of the quadriceps muscles of six random samples per group, using the RNeasy Fibrous Tissue Mini Kit (QIAGEN Gmbh, Hilden, Germany) according to the manufacturer’s instructions. RNA was stored at −80°C until the reverse transcription reaction was performed. cDNA was synthetized from 1 µg total RNA with the Omniscript RT kit (QIAGEN Gmbh, Hilden, Germany). To perform the real-time PCR reaction, cDNA was mixed with each primer (sense and antisense (0.3 µM final concentration), SYBR Green (Roche Diagnostics, Mannheim, Germany) as a fluorescent dye and H_2_O. The real-time PCR measurement of individual cDNAs was performed in triplicate using SYBR Green dye to measure duplex DNA formation with the ViiA™ 7 Real-Time PCR System (Applied Biosystems, Waltham, MA, USA). The primers sequences were designed using information contained in the public database GenBank of the National Center for Biotechnology Information (NCBI). The sequences of the primer sets used are listed in Table S1. Quantification of gene expression was performed by the method described in (54), using the 18S gene as the internal control. The amplification efficiency of each sample was calculated as described by Ramakers et al. (55).

### GSH content

GSH content was determined in superficial gastrocnemius and the oxidative part of quadriceps muscles (10 mg) using the GSH-Glo Glutathione Assay kit from Promega, following the manufacturer’s instructions.

### Western blotting

Approximately 20 mg of rat skeletal muscle (superficial gastrocnemius and the oxidative part of the quadriceps) was homogenized with a microdismembrator for 1 min at 2000 rpm (Sartorius Stedim Biotech, Aubagne, France). Muscle homogenates were then lysed on ice for 15 min with (100 µl per mg tissue) of RIPA buffer (50mM Tris-HCl pH 7.4, 150mM NaCl, 50mM NaF, 2mM EDTA, 1% NP-40, 0.5% Na-deoxycholate, 0.1% SDS, and Complete Mini protease inhibitor [Roche, Basel, Switzerland]). After lysis, the mixture was vortexed and centrifuged for 10 min at 4°C at 10,000 rpm. The supernatant was collected and the protein concentration determined using the Pierce BCA protein assay kit (ThermoFisher Scientific, Waltham, MA, USA. For each sample, 9 µg of protein was separated on a NuPAGE 4–12% Bis-Tris gel (Life technologies, Rockville, MD, USA). Proteins were electroblotted to PVDF membranes (Bio-Rad), and immunodetected using primary antibodies directed against PGC-1α (KP9803, calbiochem, 1/1000), PGC-1β (sc-373771, santa cruz, 1/1000), TFAm (ab131607, abcam, 1/1000), TOMM20 (ab78547, abcam, 1/1000), Cytochrome b (sc-11436, santa cruz, 1/1000), SOD2 (#13194, cell signaling, 1/2000), full and cleaved Caspase 3 (#9665S, cell signaling, 1/500), and Beta Actin (sc-130656, santa cruz, 1/1000). Membranes were probed with secondary antibodies conjugated to HRP directed against goat (sc-2020, santa cruz, 1/2000), rabbit (sc-2004, santa cruz, 1/2000), and mouse (sc-2055, santa cruz, 1/2000). Membranes were revealed with a chemiluminescent substrate (Clarity Western ECL substrate; Bio-Rad Laboratories, Hercules, CA, USA), and quantification was performed using the ImageJ software (National Institute of Health). Images were modified with ImageJ to remove background.

### mtDNA content

DNA was isolated from gastrocnemius and quadriceps muscles of six random samples per group, using the DNeasy Blood and Tissue Kit (QIAGEN Gmbh, Hilden, Germany) according to the manufacturer’s instructions. DNA in samples was quantified spectrophotometrically at 260 nm with a NanoDrop 2000 (ThermoFisher scientific, Waltham, USA). The DNA was subjected to real-time PCR in triplicate (10ng/µL). Relative amounts of nuclear and mitochondrial DNA were determined by comparison of amplification kinetics of Hexokinase 2 (Hk2) and COX2 (Primer sequences in Table S1, as described by (56)). mtDNA content was estimated by calculating the inverse of the CT_COX2_/CT_Hk2_ ratio.

### TUNEL assessment of apoptosis

TUNEL staining of myonuclei positive for DNA strand breaks was performed using the Click-iT® TUNEL Alexa Fluor® 647 Imaging Assay kit (ThermoFisher Scientific, Waltham, MA, USA) on randomly selected slides samples. Cross-sections (10 µm) of 3 samples per group of the glycolytic muscle gastrocnemius, and the more oxidative type vastus intermedius muscle cut with a cryostat microtome were fixed with 4% paraformaldehyde for 15 min and permeabilized with 2 mg/ml proteinase K. The TUNEL reaction mixture containing terminal deoxynucleotidyltransferase (TdT) and fluorescein-labeled dUTP was added to the sections in portions of 100 µl and then incubated for 60 min at 37°C in a humidified chamber in the dark. Section were then washed with 3% BSA and 0.1% Triton X-100 in PBS for 5 minutes, and incubated for 30 minutes at 37°C protected from light with the Click-iT^®^ Plus TUNEL reaction cocktail. After incubation, the sections were rinsed with PBS. Following embedding with ProLong diamond antifade mountant with DAPI (Life Technologies), the sections were investigated with a fluorescence microscope (x40 objective; Olympus IX83). Three to five pictures were acquired per sample. After acquisition, background noise was removed using the ImageJ software (National Institutes of Health). The number of TUNEL-stained was manually determined to avoid artefacts detection, whereas the total number of nuclei count was automated using the Icy software (v1.9.4.1) (57). Apoptotic index was calculated by counting the number of TUNEL-stained nuclei divided by the total number of nuclei multiplicated by 100.

### Statistics

Data are represented as means ± SEM. Statistical analyses were performed using unpaired *t* test or 2-way ANOVA followed by a Bonferroni’s post-test between atorvastatin treated mice in comparison to their respective controls using GraphPad Prism 5 (Graph Pad Software, Inc., San Diego, CA, USA). Statistics were then verified using RStudio (RStudio Team (2016). RStudio: Integrated Development for R. RStudio, Inc., Boston, MA URL http://www.rstudio.com/). Statistical significance is displayed as * p< 0.05: ** p < 0.01 and *** p<0.001.

## Acknowledgements

We thank the staff of the mouse facilities from Institut de Génétique et de Biologie Moléculaire et Cellulaire and Institut Clinique de la Souris from Illkirch in France. This work was supported by funds from the Centre National de la Recherche Scientifique, the Institut National de la Santé et de la Recherche Médicale, the Collège de France, the Université de Strasbourg, the Agence Nationale de la Recherche (05-PCOD-032) and by French state funds through the Agence Nationale de la Recherche ANR-10-LABX-0030-INRT under the frame programme Investissements d’Avenir labelled ANR-10-IDEX-0002-02. G.L. was supported by the Agence Nationale de la Recherche (2010BLAN1108-01). SK was supported by a grant of the Swiss National Science Foundation (31003A_156270).

## Author contribution

F.S., J.Z., D.M., B.G., S.K. and B.J. designed experiments, and F.S., U.D., A.L.C., G.L., and B.J. performed experiments and analyzed data. F.S., D.M., B.G., S.K., and J.B. wrote the manuscript.

